# Diversity, function and evolution of aquatic vertebrate genomes

**DOI:** 10.1101/2021.10.29.466026

**Authors:** Yue Song, Mengjun Yu, Suyu Zhang, Rui Zhang, Inge Seim, Xinyu Guo, Meiru Liu, Lili Yu, He Zhang, Hanbo Li, Shanshan Liu, Xin Liu, Xun Xu, Huanming Yang, Kun Wang, Shunping He, Wen Wang, Fish10K Consortium, Guangyi Fan

## Abstract

Aquatic vertebrates consist of jawed fish (cartilaginous fish and bony fish), aquatic mammals, reptiles and amphibians. Here, we present a comprehensive analysis of 630 aquatic vertebrate genomes to generate a standardized compendium of genomic data. We demonstrate its value by assessing their genome features as well as illuminating gene families related to the transition from water to land, such as *Hox* genes and olfactory receptor genes. We found that LINEs are the major transposable element (TE) type in cartilaginous fish and aquatic mammals, while DNA transposons are the dominate type in bony fish. To our surprise, TE types are not fixed in amphibians, the first group that transitioned to living on land. These results illustrate the value of a unified resource for comparative genomic analyses of aquatic vertebrates. Our data and strategy are likely to support all evolutionary and ecological research on vertebrates.

## Introduction

Aquatic vertebrates, including fish, amphibians, reptiles, and marine mammals, are ecologically, economically, and scientifically important for us, but remain understudied compared to terrestrial taxa. High-quality, chromosome-level genomes are essential for research of such species, since, finely assembled reference genomes are the basis for large-scale functional multi-omics studies. As the improvement of high-throughput sequencing technologies and analysis methods have greatly reduced the cost of price and time to sequence the genome of any species, numerous aquatic vertebrate genomes have been sequenced or under sequencing, supported by mega scientific projects such as the 10,000 Genome project (Koepfli et al., 2015; Scientists, 2009), Vertebrate Genome Project (Consortium) and the Fish10K project (Guangyi et al., 2020). Benefit from such data, we generated unified dataset and systemically evaluated genomes from aquatic vertebrates to generate a standardized compendium for comparative analyses. As results, we have provided evidences for phylogenetic controversies, distinguished genome features of major aquatic vertebrate groups, and preliminarily revealed molecular basis of the transition from aquatic to terrestrial habitats.

## Results

### Data set collection and overview

We collected *de novo* assembled genomes of 630 species from five clades of aquatic vertebrates: cartilaginous fish, bony fish, amphibians, reptiles, and marine mammals (**Supplementary Table 1**). As indicated by **Figure 1**, the majority of available genome assemblies are from fish. A unified nonredundant compendium of aquatic vertebrate genomes was created and is publicly available at https://db.cngb.org/datamart/animal/DATAani19. The resource was used to evaluate various genome wide characteristics among diverse aquatic vertebrate taxa. We assessed genome assembly quality and re-annotated TEs via a uniformed pipeline. As expected, the integrity and continuity of genome assemblies have been continuously improving for last 15 years (**Figure 2a**). We analyzed the data of jawed fishes (cartilaginous fish and bony fish) as the demonstration case, as they comprise half of extant vertebrate species and their last common ancestor with mammals can be dated back to 450 million years ago. Of 537 fish genome assemblies in our compendium, 269 (50.09%) have a scaffold N50 larger than 1 Mb and thus considered as high-quality. Among these high-quality genomes, 96 were assembled at the chromosome-level. 157 (29.23%) newly sequenced fish genomes have been obtained using stLFR linked-read sequencing technology and/or PacBio technology by the Fish10K consortium in the last two years (Bi et al., 2021; Guangyi et al., 2020; Wang et al., 2019). We reannotated the 269-species’s high-quality genome assembly using ab initio gene prediction and homology-based prediction methods, and identified around 20,000 genes for each species (See the methods section). For TEs reannotation, a uniformed pipeline was performed by s combined method of *de novo* and homology-based analysis (See the methods section), averagely 32.75% of TE content is located for these species. For comparison also reannotated the genomes of 15 amphibians, 26 reptiles, and 52 marine mammals (**Supplementary Table 2**).

**Figure 1.**
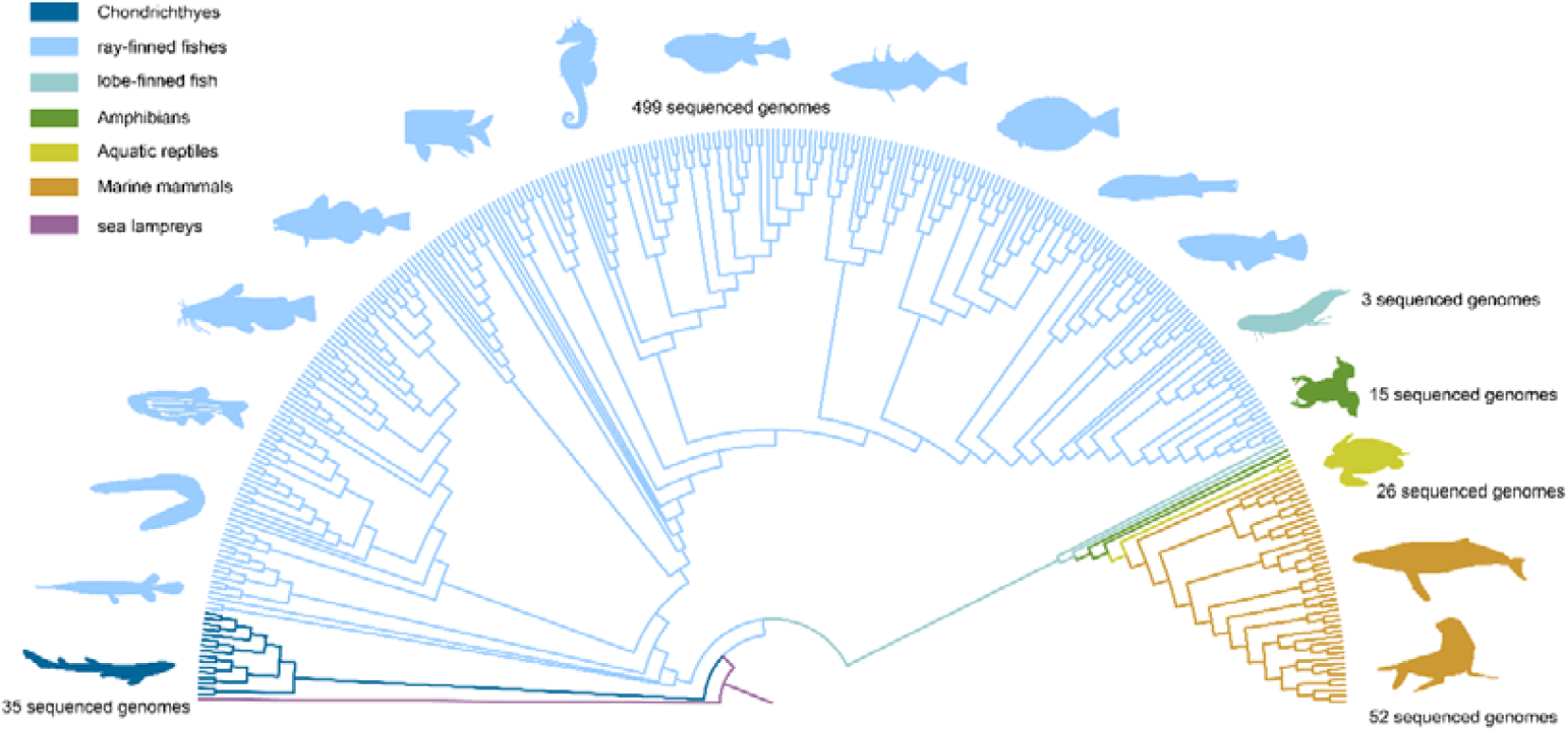
Overview of the major clades in the 630 aquatic vertebrate genome resources. Assembled nuclear genomes are shown. Note that the phylogeny is for illustrative purposes only.

**Figure 2.**
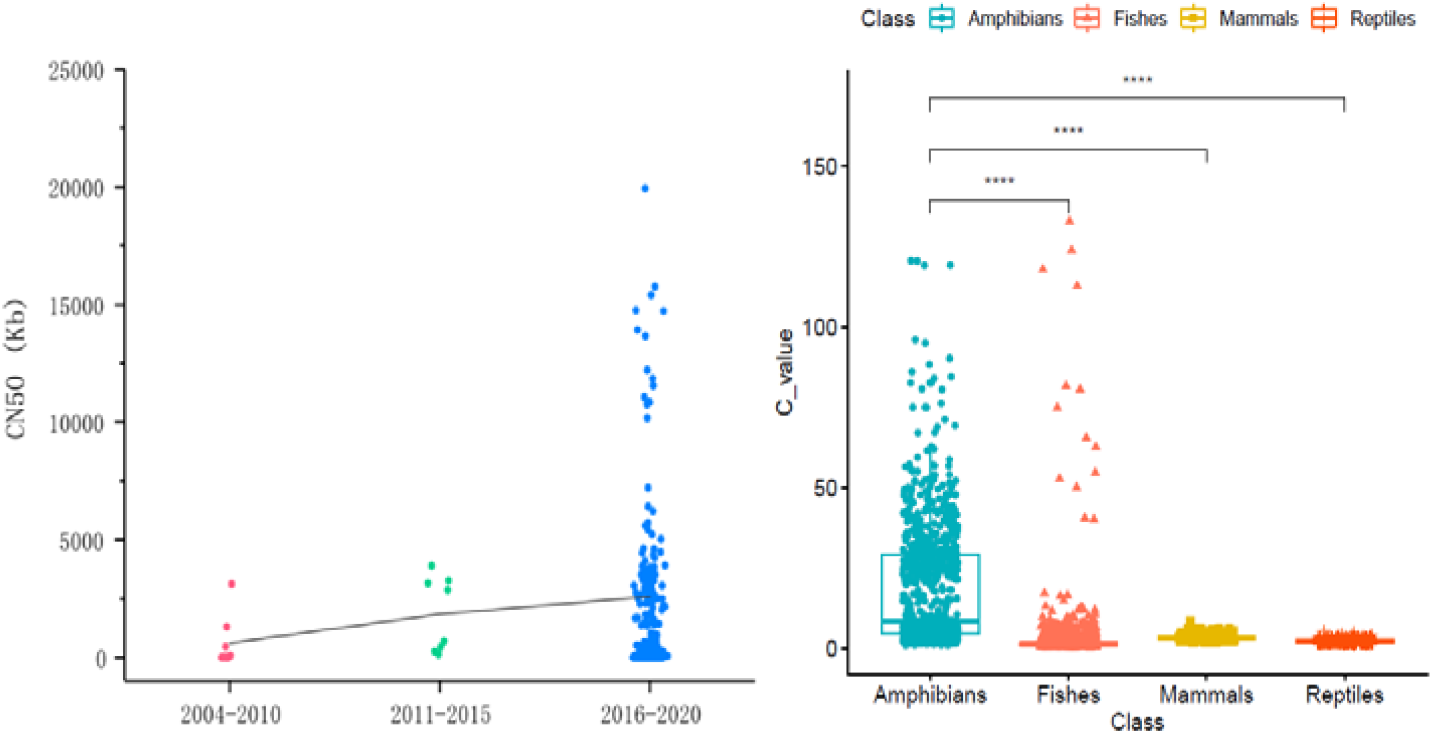
Assembly quality of the aquatic vertebrate genomes. a) Boxplot of contig N50 of 473 published genomes in fish, amphibians, aquatic reptiles and marine mammals during the period 2004-2020. Since 2015, due to the cost reduction of sequencing and the development of new sequencing technologies (e.g. Hi-C, PacBio, stLFR) (Belton et al., 2012; Rhoads and Au, 2015; Wang et al., 2019), both the integrity of genome assembly and the number of sequenced species have increased significantly in recent years. b) Boxplot of genome C-value in aquatic vertebrate groups. C-value (genome size) data was obtained from the Animal Genome Size database (Dufresne and Jeffery, 2011; Gregory, 2002a, 2005).

### Genome size and transposable elements diversity

C-value is the amount of DNA contained within a haploid nucleus (e.g., a gamete) or one half the amount in a diploid somatic cell of a eukaryotic organism, usually scaled in picograms. This method of measuring the genome size experimentally is often used to make genome-size comparisons in different species. In agreement with C-value data from the Genome Size database (Gregory, 2005) and a recent genomics study (Gregory et al., 2007; Vinogradov, 2004) (**Figure 2b**), the size of assembled genomes varied tremendously among aquatic vertebrates. Amphibians (i.e., frogs, salamanders, and cecilians) is a clade with significant variation in genome size (Liedtke et al., 2018), assembled genomes are ranged (average 4.41 Gb) from 0.5 Gb (Couch’s spadefoot toad *Scaphiopus couchii*) (Gregory, 2002b) to 32 Gb (axolotl *Ambystoma mexicanum*) (Nowoshilow et al., 2018). Although not as significant as in amphibians, genome size variations were also observed in fish (**Figure 2b**). The range in bony fish (Osteichthyes) is from 350 Mb (Tetraodontiformes) to 40 Gb (lungfishes) (Meyer et al., 2021; Wang et al., 2021), while in cartilaginous fish (Chondrichthyes) is from 3 Gb (*Chiloscyllium plagiosum*) (Zhang et al., 2020) to 6.7 Gb (cloudy catshark: *Scyliorhinus torazame*) (Hara et al., 2018).

Variations of genome size is usually related to transposable element (TE) diversity (Shao et al., 2019; Sotero-Caio et al., 2017). The average TE content of 15 amphibians, 26 reptiles, and 52 marine mammals is 40.41%, 38.20% and 34.39%, respectively (**Supplementary Table 2**). The average TE content proportion of most fishes is ∼32%. However, there are exceptions, in *Monacanthus chinensis* and *Thalassophryne amazonica*, 9.26% and 83.84% of the genome are derived from TEs, respectively (**Supplementary Table 2**). We observed an obvious inflection point between TE and genome size in bony fish (**Figure 3b**), which consistent with the relationship between TE activity and repression strategies in lungfish (Wang et al., 2021). We also observed an obvious direct, relationship between TE content and genome size in aquatic vertebrates, especially in bony fish (R^2^=0.7, p<2.2e-16) (**Figure 3b**). This relationship has been postulated to stem from major genomic rearrangements as well as missing regions stemming from limitations of DNA sequencing technologies, despite the latter can now often be resolved by long (e.g., PacBio and ONT) (Goodwin et al., 2015; Rhoads and Au, 2015) and co-barcoding sequencing technologies (e.g, stLFR and 10X genomics) (Wang et al., 2019).

**Figure 3.**
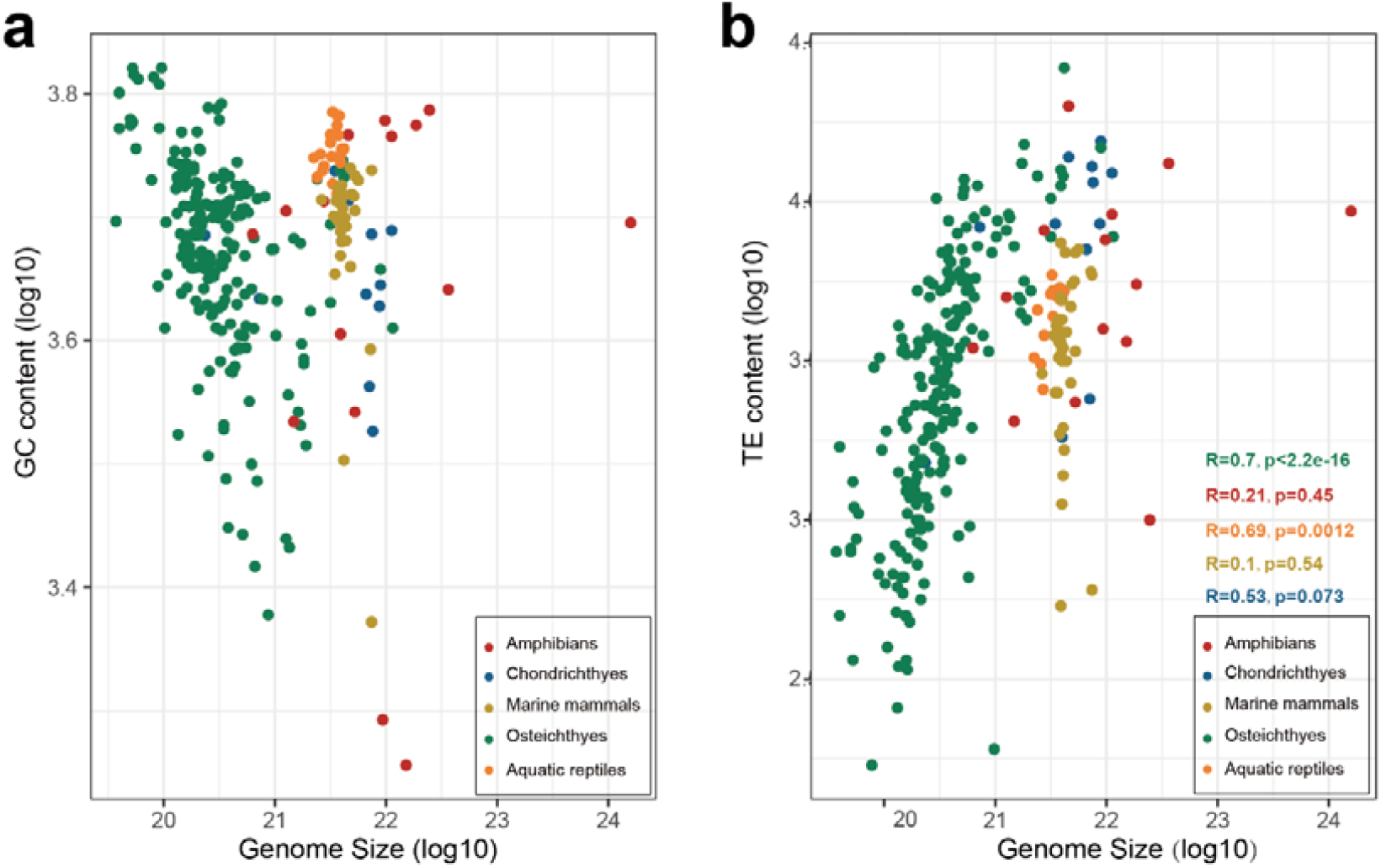
The proportional distribution of TEs in aquatic vertebrate genomes. The horizontal and vertical axis correspond to the logarithmic value of true genome size and GC content (a), TE content (b). **a)** There was no significant positive correlation between GC content and genome size; **b)** There is a positive correlation between TE and genome size, but the positive correlation is not strict in other species except Osteichthyes. This inconsistency may be related to less available genome data in different animal species.

By comparing the proportion of TEs of each genome, the proportion of TE types in species during the water-to-land evolutionary status was notably different. LINEs are the dominate component for cartilaginous fishes, and the scenario is similar for marine mammals. Meanwhile, DNA transposon is the major TE type in bony fish. The positions of the DNA transpose and LINE of Amphibians, as the evolutionary bridge from aquatic to terrestrial species, are not fixed in different species. DNA transpose is dominate in most of Anura species, but not in the three Gymnophiona species (*Geotrypetes seraphini, Rhinatrema bivittatum* and *Microcaecilia unicolor*) (**Figure 4**). For TE proportion, we proposed that one kind of special TE transformation pattern were involved in the evolutionary process from aquatic to terrestrial environment.

**Figure 4.**
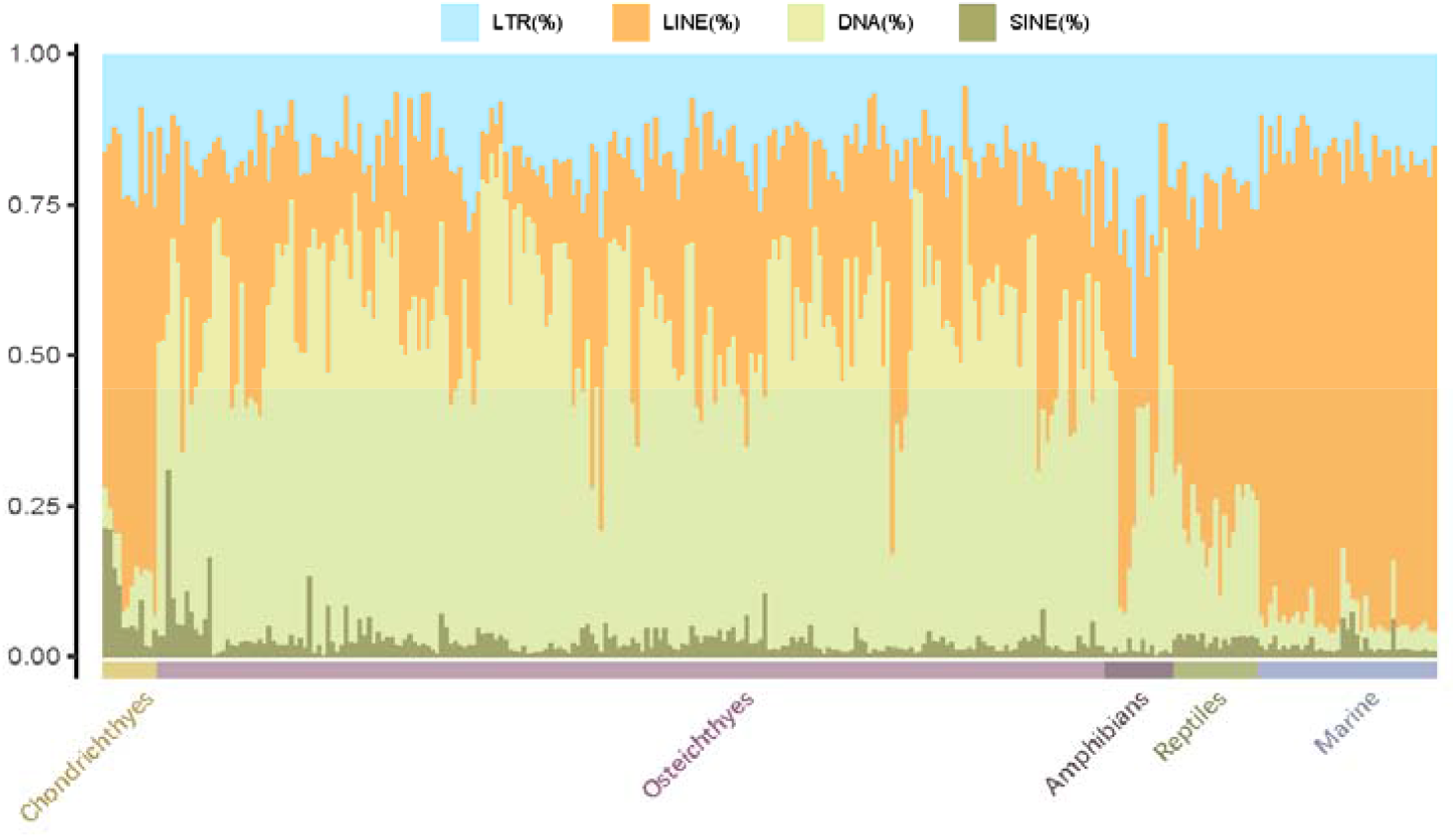
The statistics of main transposable element in marine vertebrate. LINEs in cartilaginous fishes are the dominate component (horizontal yellow bar at the bottom) and the similar scenario in marine mammals. While, DNA transposon is the top type in bony fish (horizontal purple bar at the bottom). Amphibians, as the evolutionary bridge from aquatic to terrestrial species, shown different modes of TE component in different species.

Furthermore, to detect whether there was divergence of LINE/CR1 and LINE/L2 between the Cartilaginous fish and bony fish, we constructed NJ trees with CR1 and L2 sequences from 11 bony fishes and 11 cartilaginous fishes, respectively. Although most of the CR1 sequences comes from cartilaginous fish genome and only a few of belong to bony fish, all the LINE/CR1 were clustered into two significant groups, corresponding to cartilaginous and bony fish, which is similar in LINE/L2 subtype (**Figure 5**).

**Figure 5.**
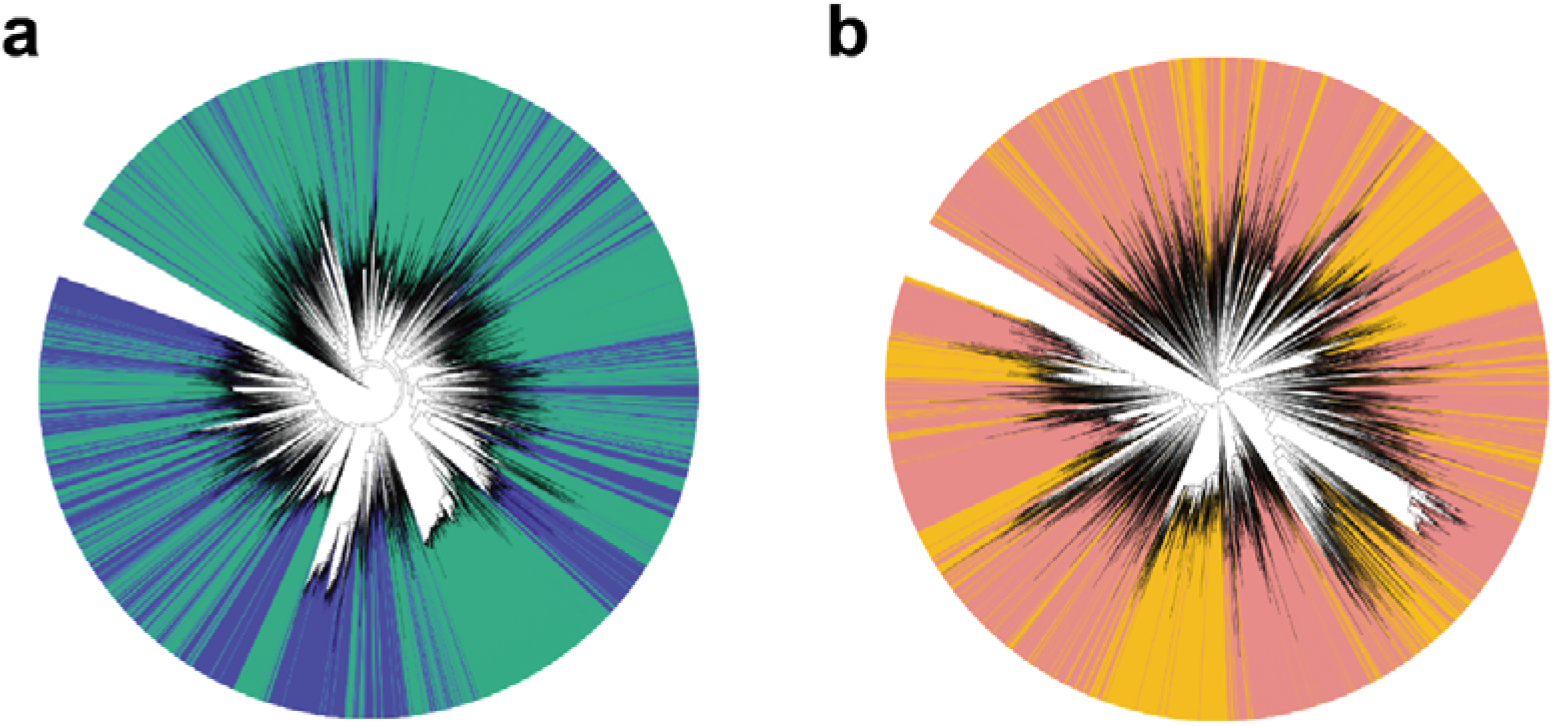
The phylogenetics trees of LINE/CR1 (a) and LINE/L2 (b). **a)** The blue branch indicated Chondrichthyes-source CR1 sequences and green branch indicated the Osteoichthys-source sequences. **b)** The orange branch indicated Chondrichthyes-source L2 sequences and pink branch indicated the Osteoichthys-source sequences. It could be observed that in (a) and (b), cartilaginous fish and bony fish have clarity difference in CR1 and L2.

### Key gene families involved in water-land transitions

In the process of evolution, great change has been taken place in marine vertebrates regardless environment or body size. To better understand such change, we highlighted *Hox* and Olfactory receptor (OR) gene families (See the methods section). *Hox* genes are belong to an important family for developmental regulation in organisms. It encodes a class of transcription factors, binding to a specific region of DNA which regulates the morphogenesis of the body axis during ontogenesis in animals, plays an important regulatory role in the development of individual embryos (Di-Poi et al., 2009; Di-Poi et al., 2010; Duboule, 1998; Lemons and McGinnis, 2006). Therefore, the study of the *Hox* gene family not only helps us understand the genetic control in animal ontogenesis, but also helps us explain the fin-to-limb transition in the evolution of vertebrates from aquatic to terrestrial at molecular level. By using published data, we identified *Hox* gene family in 63 published genomes according to the road map of evolution, the genomes are from 7 cartilaginous fishes, 47 boney fishes, 4 amphibians, 4 reptiles and 4 marine mammals. Interestingly, bony fishes are not showing the similar downward trend as observed in other species, as several WGD events happened. For example, *Salmo salar* contains 109 *Hox* genes due to its genome-wide replication events (the *Salmo*-specific WGD) (Lien et al., 2016) (**Figure 6**).

**Figure 6.**
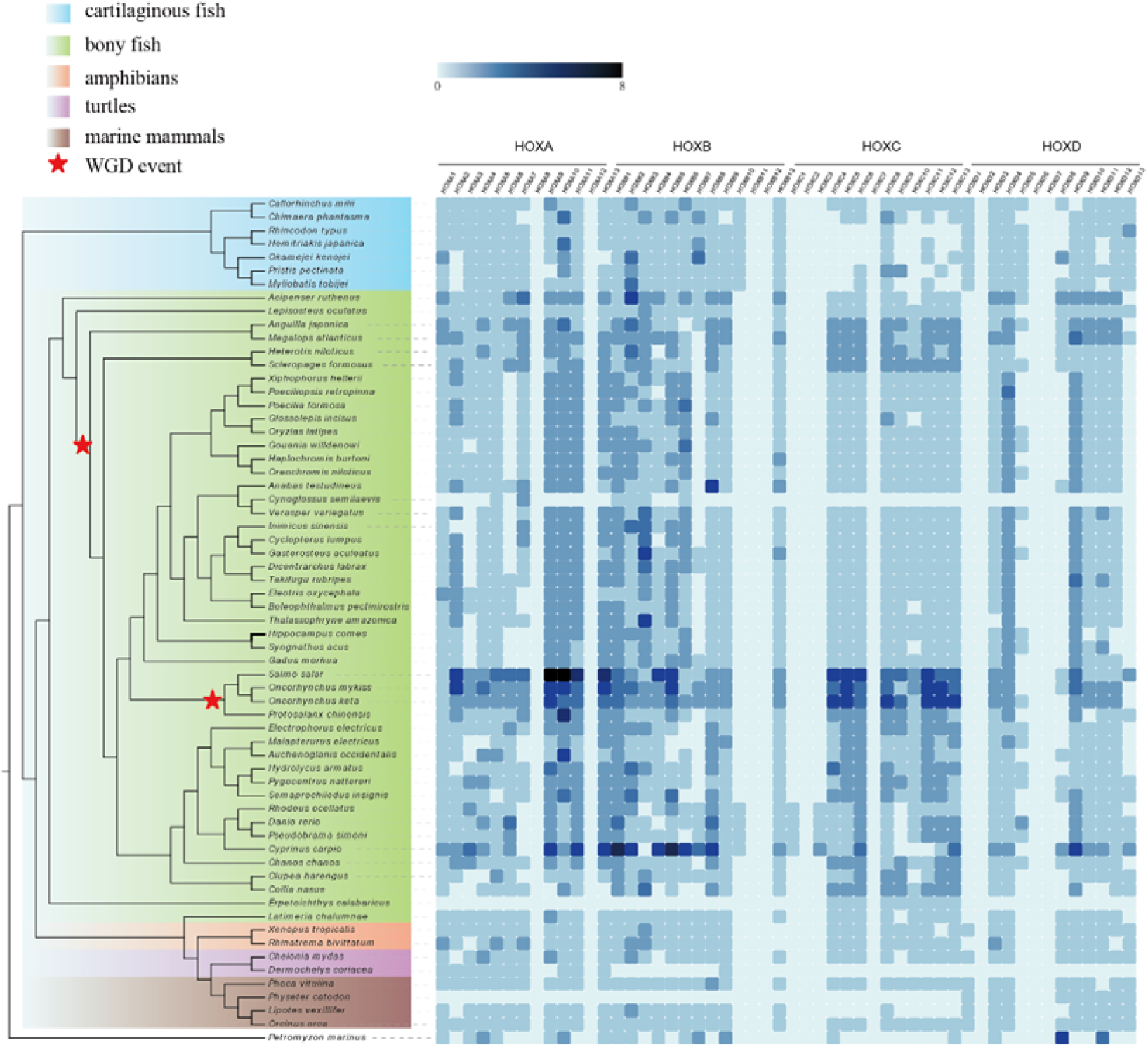
Heatmap of *Hox* gene number in different clades.

*Hox* genes are not only various in number, but also significantly different for functional gene missing. Among different cluster of *Hox* genes in these species, most of *HoxA* and *HoxB* genes are relatively conserved in different species, except for the sea lamprey. *HoxC* is low in gene number as some of such genes were missing in cartilaginous fishes and sea lampreys. Such pattern was changed dramatically in bony fish (e.g., *Acipenser ruthenus*), in which *HoxC* is remarkably increased in gene number. (**Figure 6**). This obvious change of *HoxC* gene is potentially caused by two rounds of genome-wide replication in early vertebrates (Damas et al., 2018; Venkatesh et al., 2014), as one of the duplicated homologous regions occurred a large number of gene deletions, maintained the status consistent with early vertebrate differentiation.

In previous research, vertebrate OR genes can be classified into two types, which are further classified to 11 groups (**Figure 7**, α**-**λ) corresponding to the 11 ancestral OR genes of the most common recent ancestor among vertebrate species (Freitag et al., 1998; Gaillard et al., 2004; Liu et al., 2019a; Niimura and Nei, 2005). We identified the OR genes in 62 representative aquatic and terrestrial species, where Amphibia species (such as xenopus and axolotl) possessed approximately 1000 OR genes in the genome, which is much more than other species. In Actinopterygii, the family size of OR genes are basically less than 200, normally less than 100. To our surprise, there are only less than five OR genes in each Chondrichthyes genomes in our results and few of them related to water-soluble odors detection (class δ, ε, ζand η) in cartilaginous fish. This is very different with the case of bony fish. In bony fish, the largest number of OR genes are responsible for detect odors in water. In some fish of the basal Actinopterygii, there are OR genes related to air detection (class γ). Such results suggest that the ability to use air has been developed in the early differentiation of bony fish (**Figure 7**), which consistent with previous research (Bi et al., 2021).

**Figure 7.**
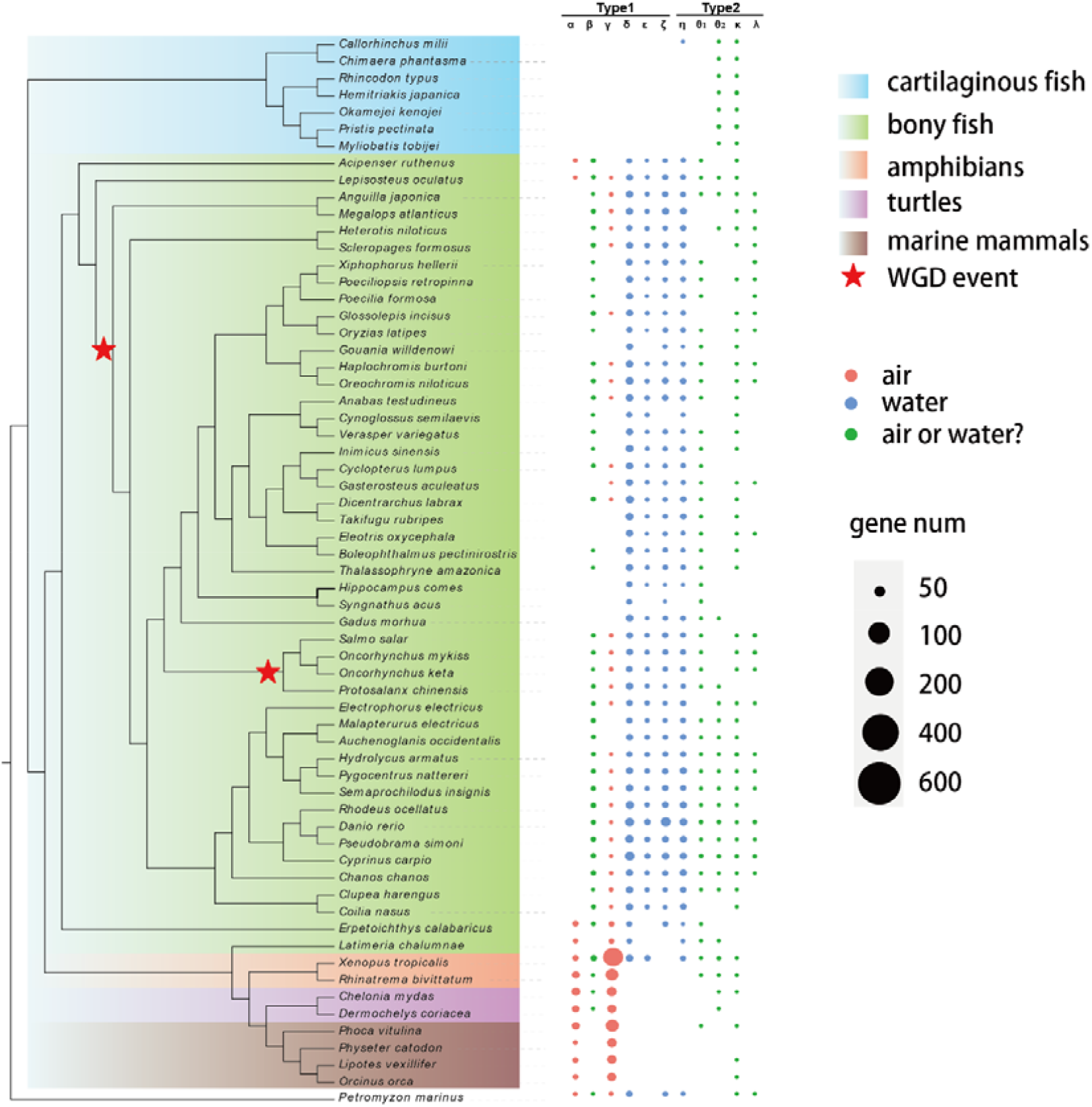
Gene number of olfactory receptors in 62 representing species, covered fish, amphibia, reptilian and marine mammals. The identified OR genes were divided into 11 groups (α-λ). “water-soluble” indicated these classes could be used to detect odors that soluble in water and “air” indicated could be used to detect odors which float through the air.

All OR genes in these typical species were used to construct phylogenetic tree. Two non-OR GPCRs were used as outgroup (NP_005292.2 and NP_037477.1) (**Figure 8**). We noticed that each branch harbors OR genes from different groups. For instance, branch (a) clustered only tetrapod OR genes (Amphibia, Aquatic reptilian and Marine mammals), so this branch may be corresponded to group α and γ. These two groups were known to present in tetrapod’s but absent in fish. To adapt both aquatic and terrestrial environments, the OR genes of Amphibia were obviously expanded across the phylogenetic tree, formed a large family to detect both water-soluble molecules and air molecules. The Sarcopterygii-specific OR genes are exhibited similar pattern of amphibians, clustered together with tetrapods OR genes (branch b), which might be involved into the early evolution from aquatic to terrestrial.

**Figure 8.**
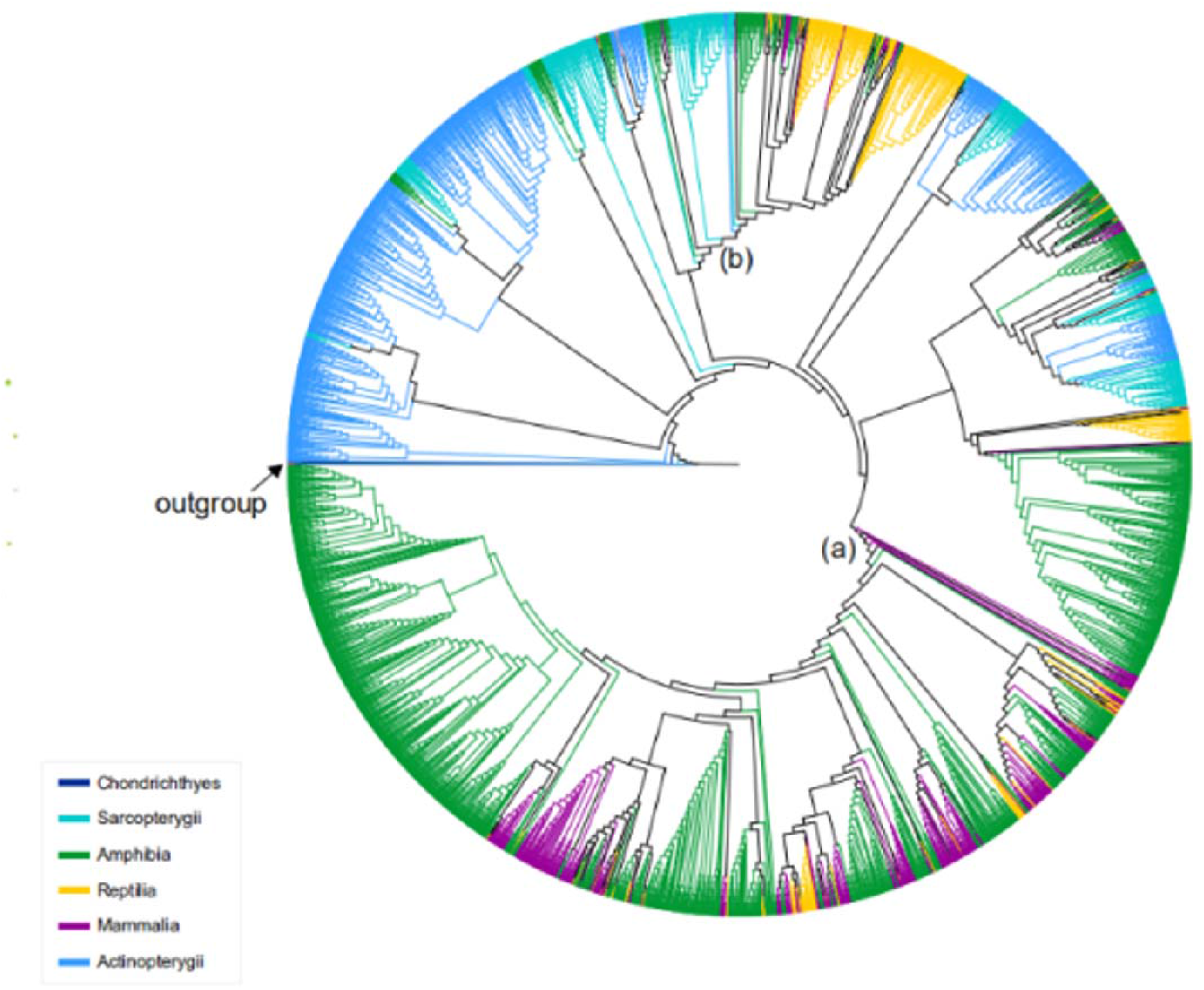
The phylogenetic tree of olfactory receptors of aquatic vertebrate genomes.

## Methods

### Genomic sequencing and assembly

To assemble the newly sequenced genome, about 150Gb (∼100X) sequence data was generated using BGI-SEQ 500 platform for each species. Then the format of sequencing data was transformed from stLFR to similar format of 10X Genomics. After the transformation, Supernova (version 2.1) (Weisenfeld et al., 2017) was used for genome assembly with default parameters. Finally, Gapcloser software (Luo et al., 2012) was used to fill the gaps within the assembly with default parameters.

### Re-annotation of transposable elements

In order to ensure the comparability of transposable elements annotation between different species, we adopted a unified method and software for transposon re-annotation. Two major types of repetitive sequences (tandem repeats and transposable elements) in this research have been reannotated. According to the method of previously research (Li et al., 2018; Liu et al., 2019b; Song et al., 2016), TRFs were identified using Tandem Repeats Finder (version 4.04) (Benson, 1999) and transposable elements (TEs) were identified by a combination of homology-based and *de novo* approaches. For homology-based approaches, different type of transposable elements sequences in Repbase (version 16.02) (Bao et al., 2015) were aligned against the assembly using RepeatMasker (version 3.2.9) (Smit et al., 2015). For *de novo* annotation, a *de novo* non-redundant repeat library was constructed using RepeatModeler (version 1.1.0.4) (Smit and Hubley, 2008) and then the newly and species-specific transposable elements were identified using RepeatMasker (Smit et al., 2015).

### Protein-coding genes annotation

Protein-coding gene were then predicted by a combination of two ways: (1) the ab initio gene prediction and (2) the homology-based annotation (Ao et al., 2015; Shao et al., 2018; Valenzano et al., 2015). For ab initio gene prediction approaches, Augustus (Stanke et al., 2006) was used with *Danio rerio* as the reference model to predicted gene models; For homology-based annotation, protein sequences of 6 related species, including zebra fish (Howe et al., 2013), spot gar (Braasch et al., 2016), elephant shark (Venkatesh et al., 2014), frog (Hellsten et al., 2010), turtle (Wang et al., 2013), dolphin (McGowen et al., 2012) and single-copy orthologs of vertebrate from BUSCO (Manni et al., 2021), were aligned against the genome assembly using BLAT software (version 0.36) (Kent, 2002) and GeneWise software (version 2.4.1) (Birney et al., 2004). The final Protein-coding gene set for each species was obtained by combining the predicted and annotated genes using Glean software (version 1.0) (Elsik et al., 2007).

### Prediction of *Hox* and olfactory receptor (OR) genes

*Hox* and olfactory receptor genes were identified using tblastn software (McGinnis and Madden, 2004) for the alignment search in each genome with homologous protein sequence of Hox and olfactory receptor genes from zebra fish, sport gar, frog and human as queries. We then predicted the structure of sequenced genes by using blast hit sequence with the software GeneWise (Birney et al., 2004), extending 2,000bp in both 3’ and 5’ directions along the genome sequences.

## References

Ao, J., Mu, Y., Xiang, L.-X., Fan, D., Feng, M., Zhang, S., Shi, Q., Zhu, L.-Y., Li, T., and Ding, Y. (2015). Genome sequencing of the perciform fish Larimichthys crocea provides insights into molecular and genetic mechanisms of stress adaptation. PLoS genetics 11, e1005118.

Bao, W., Kojima, K.K., and Kohany, O. (2015). Repbase Update, a database of repetitive elements in eukaryotic genomes. Mob DNA 6, 11.

Belton, J.-M., McCord, R.P., Gibcus, J.H., Naumova, N., Zhan, Y., and Dekker, J. (2012). Hi–C: a comprehensive technique to capture the conformation of genomes. Methods 58, 268–276.

Benson, G. (1999). Tandem repeats finder: a program to analyze DNA sequences. Nucleic acids research 27, 573–580.

Bi, X., Wang, K., Yang, L., Pan, H., Jiang, H., Wei, Q., Fang, M., Yu, H., Zhu, C., Cai, Y., et al. (2021). Tracing the genetic footprints of vertebrate landing in non-teleost ray-finned fishes. Cell 184, 1377–1391 e1314.

Birney, E., Clamp, M., and Durbin, R. (2004). GeneWise and genomewise. Genome research 14, 988–995.

Braasch, I., Gehrke, A.R., Smith, J.J., Kawasaki, K., Manousaki, T., Pasquier, J., Amores, A., Desvignes, T., Batzel, P., and Catchen, J.M. (2016). The spotted gar genome illuminates vertebrate evolution and facilitates human-teleost comparisons. Nature Genetics 48, 427–437.

Consortium, G.K. Vertebrate Genomes Project (VGP).

Damas, J., Kim, J., Farre, M., Griffin, D.K., and Larkin, D.M. (2018). Reconstruction of avian ancestral karyotypes reveals differences in the evolutionary history of macro-and microchromosomes. Genome Biol 19, 155.

Di-Poi, N., Montoya-Burgos, J.I., and Duboule, D. (2009). Atypical relaxation of structural constraints in Hox gene clusters of the green anole lizard. Genome Res 19, 602–610.

Di-Poi, N., Montoya-Burgos, J.I., Miller, H., Pourquie, O., Milinkovitch, M.C., and Duboule, D. (2010). Changes in Hox genes’ structure and function during the evolution of the squamate body plan. Nature 464, 99–103.

Duboule, D. (1998). Vertebrate hox gene regulation: clustering and/or colinearity? Current opinion in genetics & development 8, 514–518.

Dufresne, F., and Jeffery, N. (2011). A guided tour of large genome size in animals: what we know and where we are heading. Chromosome Res 19, 925–938.

Elsik, C.G., Mackey, A.J., Reese, J.T., Milshina, N.V., Roos, D.S., and Weinstock, G.M. (2007). Creating a honey bee consensus gene set. Genome biology 8, R13.

Freitag, J., Ludwig, G., Andreini, I., Rössler, P., and Breer, H. (1998). Olfactory receptors in aquatic and terrestrial vertebrates. Journal of Comparative Physiology A 183, 635–650.

Gaillard, I., Rouquier, S., and Giorgi, D. (2004). Olfactory receptors. Cellular and Molecular Life Sciences CMLS 61, 456–469.

Goodwin, S., Gurtowski, J., Ethe-Sayers, S., Deshpande, P., Schatz, M.C., and McCombie, W.R. (2015). Oxford Nanopore sequencing, hybrid error correction, and de novo assembly of a eukaryotic genome. Genome research 25, 1750–1756.

Gregory, T.R. (2002a). Animal genome size database. http://www.genomesizecom.

Gregory, T.R. (2002b). Genome size and developmental complexity. Genetica 115, 131–146.

Gregory, T.R. (2005). Genome size evolution in animals. In The evolution of the genome (Elsevier), pp. 3–87.

Gregory, T.R., Nicol, J.A., Tamm, H., Kullman, B., Kullman, K., Leitch, I.J., Murray, B.G., Kapraun, D.F., Greilhuber, J., and Bennett, M.D. (2007). Eukaryotic genome size databases. Nucleic Acids Res 35, D332–338.

Guangyi, F., Yue, S., Liandong, Y., Xixoyun, H., Suyu, Z., Mengqi, Z., Xianwei, Y., Yue, C., Yongxin, L., Shanshan, L., et al. (2020). Fish10K Pilot Project Data GigaScience Database. GigaScience Database.

Hara, Y., Yamaguchi, K., Onimaru, K., Kadota, M., Koyanagi, M., Keeley, S.D., Tatsumi, K., Tanaka, K., Motone, F., and Kageyama, Y. (2018). Shark genomes provide insights into elasmobranch evolution and the origin of vertebrates. Nature ecology & evolution 2, 1761–1771.

Hellsten, U., Harland, R.M., Gilchrist, M.J., Hendrix, D., Jurka, J., Kapitonov, V., Ovcharenko, I., Putnam, N.H., Shu, S., and Taher, L. (2010). The genome of the Western clawed frog Xenopus tropicalis. Science 328, 633–636.

Howe, K., Clark, M.D., Torroja, C.F., Torrance, J., Berthelot, C., Muffato, M., Collins, J.E., Humphray, S., McLaren, K., and Matthews, L. (2013). The zebrafish reference genome sequence and its relationship to the human genome. Nature 496, 498.

Kent, W.J. (2002). BLAT—the BLAST-like alignment tool. Genome research 12, 656–664.

Koepfli, K.P., Paten, B., Genome, K.C.o.S., and O’Brien, S.J. (2015). The Genome 10K Project: a way forward. Annu Rev Anim Biosci 3, 57–111.

Lemons, D., and McGinnis, W. (2006). Genomic evolution of Hox gene clusters. Science 313, 1918–1922.

Li, C., Liu, X., Liu, B., Ma, B., Liu, F., Liu, G., Shi, Q., and Wang, C. (2018). Draft genome of the Peruvian scallop Argopecten purpuratus. GigaScience 7, giy031.

Liedtke, H.C., Gower, D.J., Wilkinson, M., and Gomez-Mestre, I. (2018). Macroevolutionary shift in the size of amphibian genomes and the role of life history and climate. Nat Ecol Evol 2, 1792–1799.

Lien, S., Koop, B.F., Sandve, S.R., Miller, J.R., Kent, M.P., Nome, T., Hvidsten, T.R., Leong, J.S., Minkley, D.R., and Zimin, A. (2016). The Atlantic salmon genome provides insights into rediploidization. Nature 533, 200–205.

Liu, A., He, F., Shen, L., Liu, R., Wang, Z., and Zhou, J. (2019a). Convergent degeneration of olfactory receptor gene repertoires in marine mammals. BMC Genomics 20, 977.

Liu, H.-P., Xiao, S.-J., Wu, N., Wang, D., Liu, Y.-C., Zhou, C.-W., Liu, Q.-Y., Yang, R.-B., Jiang, W.-K., and Liang, Q.-Q. (2019b). The sequence and de novo assembly of Oxygymnocypris stewartii genome. Scientific data 6, 190009.

Luo, R., Liu, B., Xie, Y., Li, Z., Huang, W., Yuan, J., He, G., Chen, Y., Pan, Q., and Liu, Y. (2012). SOAPdenovo2: an empirically improved memory-efficient short-read de novo assembler. GigaScience 1, 18.

Manni, M., Berkeley, M.R., Seppey, M., Simao, F.A., and Zdobnov, E.M. (2021). BUSCO update: novel and streamlined workflows along with broader and deeper phylogenetic coverage for scoring of eukaryotic, prokaryotic, and viral genomes. arXiv preprint arXiv:210611799.

McGinnis, S., and Madden, T.L. (2004). BLAST: at the core of a powerful and diverse set of sequence analysis tools. Nucleic acids research 32, W20–W25.

McGowen, M.R., Grossman, L.I., and Wildman, D.E. (2012). Dolphin genome provides evidence for adaptive evolution of nervous system genes and a molecular rate slowdown. Proceedings of the Royal Society B: Biological Sciences 279, 3643–3651.

Meyer, A., Schloissnig, S., Franchini, P., Du, K., Woltering, J.M., Irisarri, I., Wong, W.Y., Nowoshilow, S., Kneitz, S., Kawaguchi, A., et al. (2021). Giant lungfish genome elucidates the conquest of land by vertebrates. Nature 590, 284–289.

Niimura, Y., and Nei, M. (2005). Evolutionary dynamics of olfactory receptor genes in fishes and tetrapods. Proceedings of the National Academy of Sciences 102, 6039–6044.

Rhoads, A., and Au, K.F. (2015). PacBio sequencing and its applications. Genomics, proteomics & bioinformatics 13, 278–289.

Scientists, G.K.C.o. (2009). Genome 10K: a proposal to obtain whole-genome sequence for 10 000 vertebrate species. Journal of Heredity 100, 659–674.

Shao, C., Li, C., Wang, N., Qin, Y., Xu, W., Liu, Q., Zhou, Q., Zhao, Y., Li, X., Liu, S., et al. (2018). Chromosome-level genome assembly of the spotted sea bass, Lateolabrax maculatus. GigaScience 7.

Shao, F., Han, M., and Peng, Z. (2019). Evolution and diversity of transposable elements in fish genomes. Sci Rep 9, 15399.

Smit, A., and Hubley, R. (2008). RepeatModeler Open-1.0. Available fom http://www.repeatmaskerorg.

Smit, A., Hubley, R., and Green, P. (2015). RepeatMasker Open-4.0. 2013–2015.

Song, L., Bian, C., Luo, Y., Wang, L., You, X., Li, J., Qiu, Y., Ma, X., Zhu, Z., and Ma, L. (2016). Draft genome of the Chinese mitten crab, Eriocheir sinensis. GigaScience 5, 5.

Sotero-Caio, C.G., Platt, R.N., 2nd, Suh, A., and Ray, D.A. (2017). Evolution and Diversity of Transposable Elements in Vertebrate Genomes. Genome Biol Evol 9, 161–177.

Stanke, M., Keller, O., Gunduz, I., Hayes, A., Waack, S., and Morgenstern, B. (2006). AUGUSTUS: ab initio prediction of alternative transcripts. Nucleic acids research 34, W435–W439.

Valenzano, D.R., Benayoun, B.A., Singh, P.P., Zhang, E., Etter, P.D., Hu, C.-K., Clément-Ziza, M., Willemsen, D., Cui, R., and Harel, I. (2015). The African turquoise killifish genome provides insights into evolution and genetic architecture of lifespan. Cell 163, 1539–1554.

Venkatesh, B., Lee, A.P., Ravi, V., Maurya, A.K., Lian, M.M., Swann, J.B., Ohta, Y., Flajnik, M.F., Sutoh, Y., and Kasahara, M.J.N. (2014). Elephant shark genome provides unique insights into gnathostome evolution. 505, 174.

Vinogradov, A.E. (2004). Genome size and extinction risk in vertebrates. Proceedings of the Royal Society of London Series B: Biological Sciences 271, 1701–1705.

Wang, K., Wang, J., Zhu, C., Yang, L., Ren, Y., Ruan, J., Fan, G., Hu, J., Xu, W., Bi, X., et al. (2021). African lungfish genome sheds light on the vertebrate water-to-land transition. Cell 184, 1362–1376 e1318.

Wang, O., Chin, R., Cheng, X., Wu, M.K.Y., Mao, Q., Tang, J., Sun, Y., Anderson, E., Lam, H.K., Chen, D., et al. (2019). Efficient and unique cobarcoding of second-generation sequencing reads from long DNA molecules enabling cost-effective and accurate sequencing, haplotyping, and de novo assembly. Genome Res 29, 798–808.

Wang, Z., Pascual-Anaya, J., Zadissa, A., Li, W., Niimura, Y., Huang, Z., Li, C., White, S., Xiong, Z., Fang, D., et al. (2013). The draft genomes of soft-shell turtle and green sea turtle yield insights into the development and evolution of the turtle-specific body plan. Nat Genet 45, 701–706.

Weisenfeld, N.I., Kumar, V., Shah, P., Church, D.M., and Jaffe, D.B. (2017). Direct determination of diploid genome sequences. Genome research 27, 757–767.

Zhang, Y., Gao, H., Li, H., Guo, J., Ouyang, B., Wang, M., Xu, Q., Wang, J., Lv, M., and Guo, X. (2020). The white-spotted bamboo shark genome reveals chromosome rearrangements and fast-evolving immune genes of cartilaginous fish. Iscience 23, 101754.

